# Assessing sequence-based protein-protein interaction predictors for use in therapeutic peptide engineering

**DOI:** 10.1101/2021.09.29.462289

**Authors:** François Charih, Kyle K. Biggar, James R. Green

## Abstract

Engineering peptides to achieve a desired therapeutic effect through the inhibition of a specific target activity or protein interaction is a non-trivial task. Few of the existing *in silico* peptide design algorithms generate target-specific peptides. Instead, many methods produce peptides that achieve a desired effect through an unknown mechanism. In contrast with resource-intensive high-throughput experiments, *in silico* screening is a cost-effective alternative that can prune the space of candidates when engineering target-specific peptides. Using a set of FDA-approved peptides we curated specifically for this task, we assess the applicability of several sequence-based protein-protein interaction predictors as a screening tool within the context of peptide therapeutic engineering. We show that similarity-based protein-protein interaction predictors are more suitable for this purpose than the state-of-the-art deep learning methods publicly available at the time of writing. We also show that this approach is mostly useful when designing new peptides against targets for which naturally-occurring interactors are already known, and that deploying it for *de novo* peptide engineering tasks may require gathering additional target-specific training data. Taken together, this work offers evidence that supports the use of similarity-based protein-protein interaction predictors for peptide therapeutic engineering, especially peptide analogs.

## Introduction

### In silico screening for peptide therapeutic engineering

Peptide therapeutics represent a unique opportunity to interfere with abnormal enzymatic activity or to disrupt protein-protein interactions (PPIs) in a targeted fashion. In contrast with small molecules, peptides can often be designed for very high specificity against their target, thereby conferring them an advantageous safety profile. Given that peptides are simply chains of amino acids, the chemical space they span is, for all intents and purposes, chemically accessible in its entirety.

In spite of their advantages, peptide therapeutics represent an exceedingly small fraction of all approved therapeutics, with only 70 compounds approved in the U.S., Europe and Japan as of 2020^1^. In practice, oral delivery of peptide therapeutics remains a sizable challenge due to their low bioavailability caused by the presence of proteases in the gastrointestinal tract, and size and charge constraints associated with the permeability of the mucosal membrane of the gut, among others^2^. Recently, Semaglutide, which was approved by the FDA for the treatment of diabetes and obesity, became the first oral Glucagon-like peptide receptor peptide therapy approved in the US.

As a result, almost all peptides currently in clinical use must be delivered intravenously^3^. Nevertheless, there is a shared optimism that advances in targeted delivery technologies will significantly lower the barriers associated with oral peptide delivery^3^. For example, liposomal delivery of peptide therapeutics appears to be promising option for multiple routes of administration (oral, intranasal, pulmonary, *etc*.)^4^. Other technologies such as hydrogels and permeation enhancing molecules are also currently being investigated, though more work needs to be done before they can be brought to the clinic^3^. It is thus reasonably foreseeable that peptides may garner a renewed interest in the near future.

*In silico* peptide engineering for therapeutic applications is already an active area of research, with multiple groups having leveraged advances in deep learning to develop novel algorithms for the development of antimicrobial peptides^5–7^, anti-cancer peptides^8^, and anti-hypertensive peptides^9^, among others. Many of these methods generate peptides expected to achieve a certain effect; however, their exact target is often unknown. Some of these methods leverage neural networks trained on large datasets to learn the distribution of peptides possessing the desired properties. Peptides can then be sampled from such distribution to generate novel peptides that are similar, but different to those in the training set^5, 8^. Other peptide engineering methods evaluate the fitness of peptides in *vitro* (e.g. IC_50_ in^6^).

Surprisingly few target-specific peptide engineering approaches have been developed^10, 11^. This is unsurprising, given that validated protein-peptide interactions are relatively scarce in comparison with protein-protein interactions. As such, it is difficult to develop general-purpose peptide engineering algorithms that could be broadly applied to design a peptide that interacts with one’s protein target of choice. To the best of our knowledge, InSiPS ^11^ is the only entirely *in silico* method that addresses this problem as of today.

Regardless of the peptide engineering approach used, the ability to identify a therapeutic peptide’s target or to evaluate the likelihood that it will interact specifically with a target protein is highly desirable. Such a method would provide a quick, cheap way to identify candidates in preparation for more time-consuming and expensive *in vitro* or *in vivo* validation experiments.

### Sequence-based protein-protein interaction predictors

Several sequence-based PPI prediction algorithms can evaluate the likelihood of an interaction between proteins. However, no widely used method has been developed specifically for the prediction of protein-peptide interactions. In theory, nothing precludes traditional PPI predictors from making predictions for pairs involving a short peptide, but in practice many of these predictors are trained on pairs involving large proteins. As such, it is not yet clear how applicable these methods are for protein-peptide interactions.

Though generally thought to provide less accurate predictions than structure-based methods, sequence-based predictors are widely used and appreciated because they can generate predictions for any pair of proteins, so as long as an amino acid sequence is available; however, it should be mentioned that several predictors do require proteins to have a minimum length. While protein-peptide docking simulations can provide a wealth of thermodynamic information that sequence-based predictors cannot, few proteins are amenable to these experiments. In general, the resolution of protein structures required for these simulations should be in the sub-2.5Å range^12, 13^, of which there are relatively few, though reasonably accurate structures predicted with AlphaFold 2^14^ are now available for most proteins. Furthermore, sequence-based predictors tend to require far fewer computational resources to run compared with structure-based methods which may require iterative docking, a resource-intensive procedure. In fact, our massively parallel implementation of the SPRINT algorithm^15^ can predict the entire human interactome in under an hour using a 40-core machine with 64 GB memory. As such, sequence-based PPI predictors provide a unique opportunity to quickly and inexpensively screen synthetically designed peptides and to assess their potential as candidates in wet lab experiments.

Two broad types of approaches are currently being used to predict protein-protein interactions: similarity-based methods and machine learning-based methods (Figure 1). Similarity-based methods such as PIPE4^16^ and SPRINT^15^ score proteins based on the fundamental idea that a pair of known interacting proteins P1 and P2 provides evidence for an interaction between the query proteins *Q*1 and *Q*2 if *P*1 is similar to *Q*1 and *P*2 is similar to *Q*2. These methods, in essence, quantify the strength of the evidence for an interaction under this assumption, using substitution matrices such as PAM120 or BLOSUM64 to assess the similarity between query and interacting protein pairs. These methods are close to what one may call *instance-based learning* in machine learning terminology. In contrast, machine learning-based methods “learn” to recognize patterns or features that occur frequently in interacting proteins. To make predictions, the machine learning models then look for the presence or extent of these patterns in query protein pairs. Depending on the authors, the predictors may learn patterns from the physicochemical properties of the protein sequences or simply from the amino acid composition (eg. di-and tri-peptide composition, pseudo-amino acid composition, etc.). The most recent models tend to implement the latter approach with powerful deep learning models to learn the “grammar” of interacting proteins^17–19^.

**Figure 1.**
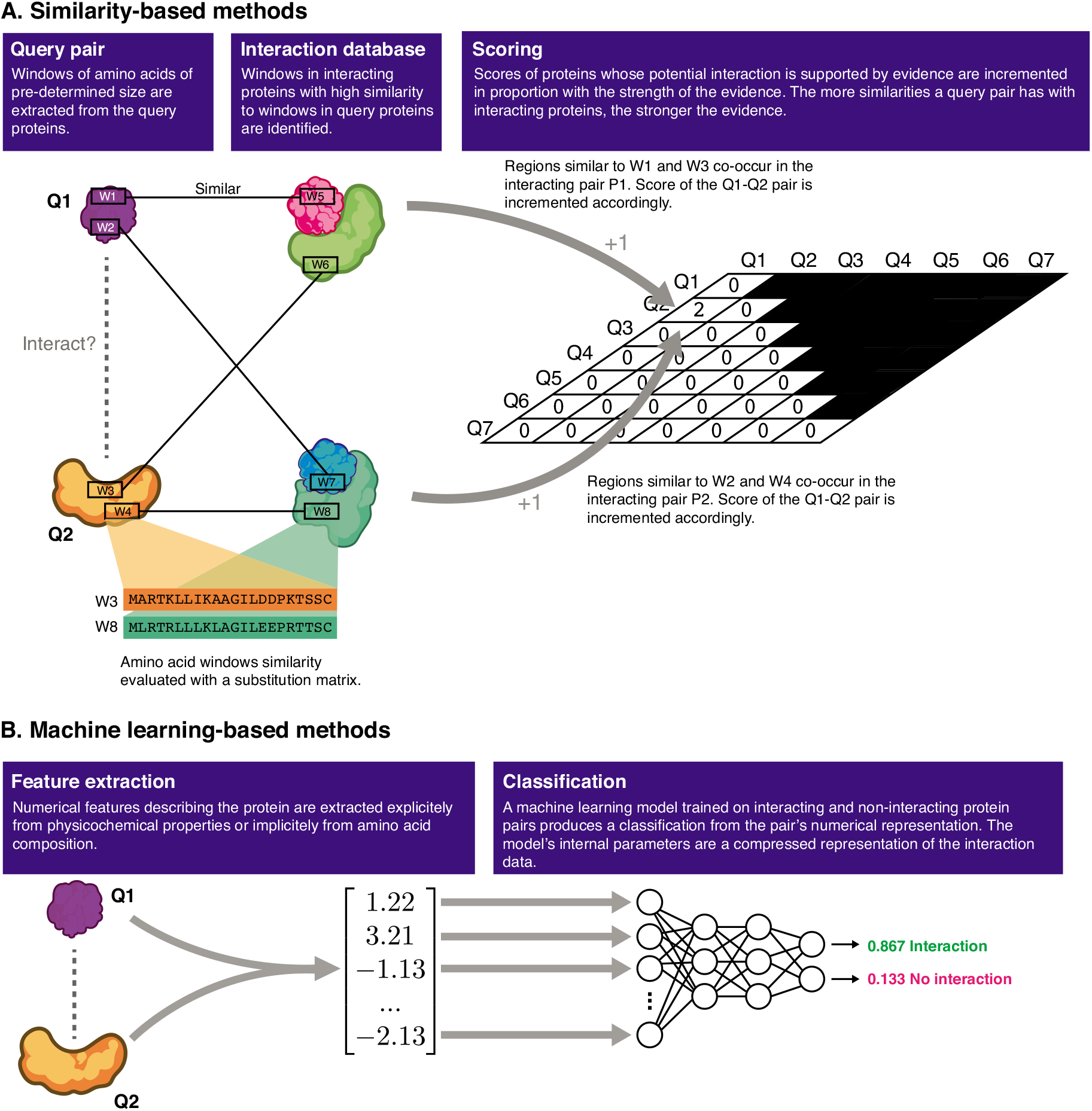
Similarity-and machine learning-based protein-protein interaction predictors. Similarity-based methods differ from machine learning-based methods in that the evidence supporting an interaction is quantified by *counting* the number of similarities to known instances of interacting proteins in a database. Machine learning-based methods are *trained* to learn patterns found in interacting protein pairs, and identify those patterns in the query proteins.

The PIPE sequence-based PPI predictor^16^ has been successfully used to design novel peptides that interact with a selected target. The *In Silico* Peptide Synthesizer (InSiPS)^11^, built around PIPE, uses genetic algorithms to explore the peptide space and maximize peptide fitness. The fitness function is designed such that the predicted interaction score between the peptide and the target is maximized, but the interaction score between the peptide and all other proteins is minimized. This ensures that the algorithm favours peptides that are predicted to interact specifically with the protein target. So far, InSiPS has been demonstrated to produce valid peptides, but no published work has shown that it can produce valid peptide binding to human targets. Furthermore, little work has been done to validate the use of recent deep learning PPI prediction models for applications in peptide engineering.

### The one-to-all curve for a qualitative assessment of peptide quality

The one-to-all curve is a simple, yet informative visualization of the interaction landscape of a protein-binding compound. Information contained within the curve has been shown to be useful in the context of PPI^20^ and miRNA target prediction tasks^21^. It is obtained by plotting the predicted interaction score of *all* proteins expressed in the target organism, tissue, or subcellular location as a function of the ranking of that score, relative to the score of other proteins, *for one peptide*. The morphology of the curve enables a qualitative assessment of a peptide in terms of its ability to bind the intended target and its specificity.

Within the context of peptide-protein interactions, most one-to-all curves for peptides are L-shaped or sigmoidal. In the more common L-shaped curve, most proteins lie along a baseline representing the proteins to which a very low interaction score with the peptide is predicted, indicating that an interaction is not supported by the evidence (training data) used to train the model. In the context of peptide engineering, all but very few peptides will produce a curve where the targeted protein lies along the baseline (Figure 2A), consistent with the idea that very few peptides will actually bind to a given protein target.

**Figure 2.**
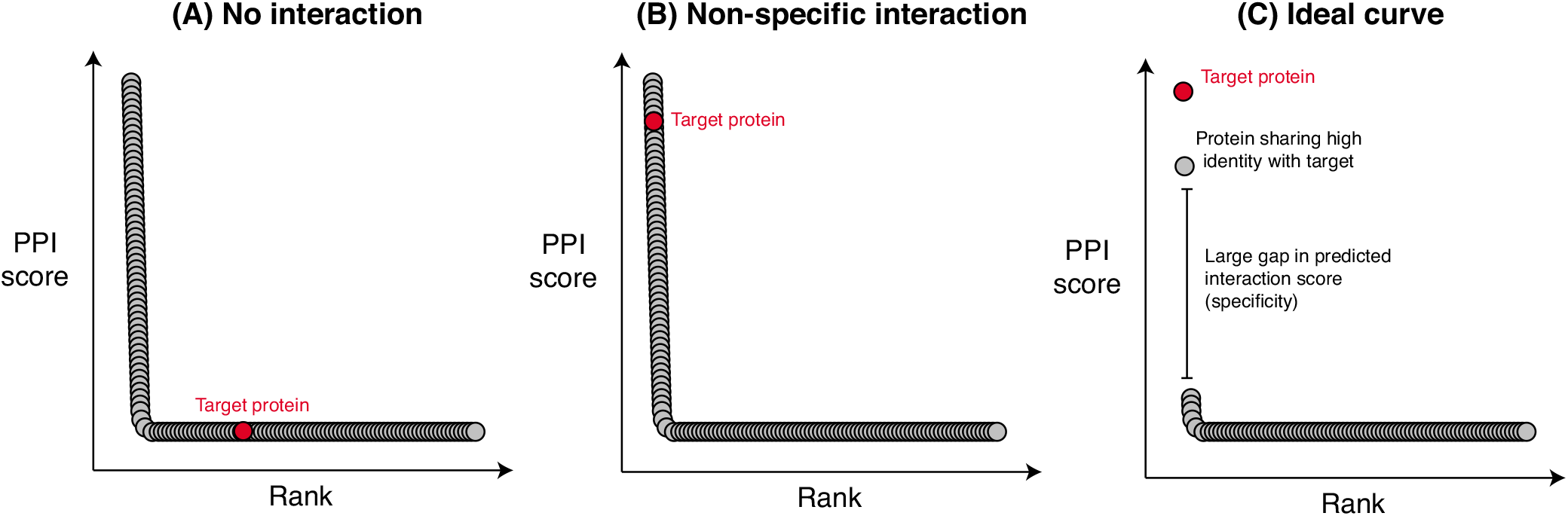
Representative one-to-all curves. Many one-to-all curves adopt one of three morphologies: (A) no interaction between the peptide and the target, (B) non-specific interaction where the peptide is predicted to interact with many proteins in addition to the target, and (C) where the peptide specifically interacts with the intended protein target (and possibly, closely related proteins).

The number of proteins which lie above the elbow of the curve provide information with respect to the specificity of the peptide, i.e. whether the peptide is predicted to interact with multiple proteins. As such, it is not only desirable for the target protein to be as high above the elbow as possible, as it must be one of the few high-scoring proteins, it must also be higher scoring than other non-target proteins. In many cases, a highly non-specific interaction with a protein target will be predicted, yielding a curve such as the one shown in Figure 2B. Such peptides are unlikely to be useful in practice, due to the high likelihood of off-target interactions, potentially leading to undesirable side effects.

Figure 2C represents the most desirable scenario for a good peptide therapeutic candidate. The target protein is strongly predicted to interact with the target protein, and the only other proteins achieving a similar score are proteins with high similarity to the target (isoforms, for example). In such cases, peptide specificity may be further optimized *in vitro*.

The y axis of the one-to-all curve is not always easy to interpret, particularly when the PPI prediction score is unbounded. Most machine learning-based predictors convert raw prediction scores into probabilities between 0 and 1 using the *softmax* function. However, similarity-based methods instead output a number that reflects the strength of the evidence, as is the case with SPRINT^15^. In such cases, it is the user’s responsibility to identify an appropriate threshold value on the score above which protein pairs are predicted to interact. This can be done using cross-validation experiments.

### Contributions

Herein, we evaluate the applicability of sequence-based PPI predictors in identifying peptides fit for *in vitro* or *in vivo* validation. Using a set of FDA-approved peptides, we compare three state-of-the-art sequence-based PPI predictors and their usefulness in evaluating peptides for creation of peptide analogous to endogenously expressed proteins, and for *de novo* peptide design for novel protein targets. We believe that this work has significant implications for peptide engineering and could guide use of PPI predictors in future peptide engineering pipelines.

## Methods

The code and instructions needed to reproduce the data and analyses presented here are available in the GitHub repository: https://github.com/GreenCUBIC/PeptideScreening.

### Assumption

Several peptides have been approved by the FDA for the treatment of a number of conditions such as heart failure and diabetes. Given that these peptides have been shown to be safe, one could deduce that these peptides interact specifically with their target, i.e. interact with few proteins other than their intended target. Off-target interactions are unlikely to be numerous, as this would likely cause unintended side effects.

This assumption underpins the interpretation of one-to-all curves generated as part of this work. In other words, FDA-approved peptides should, in theory, rank their known target protein among the first proteins on the one-to-all curve and score much higher than other proteins.

### Curating FDA-approved peptides

To identify suitable peptide therapeutics for assessment with PPI predictors, we retrieved a list of all therapeutic peptides approved by the FDA for therapeutic use as of 2017, as compiled previously^22^. Of those 69 peptides, we selected those whose length was greater than 20 amino acids and only comprised standard amino acids (see Table 1), leaving a total of 13 peptides. This additional round of curation was necessary, because certain predictors cannot predict interactions involving very small protein fragments and/or non-standard amino acids. Finally, for each peptide, we retrieved the associated protein target(s) from the DrugBank database^23^.

**Table 1.** FDA-approved peptides evaluated in this study

| Compound Name | Molecular Target | Chemical Basis | Length | Known Targets (Accession ID) |
| --- | --- | --- | --- | --- |
| Corticotropin | MC receptors | native | 39 | Q01726;Q01718;P41968;P32245;P33032 |
| Calcitonin (salmon) | Calcitonin receptor | native | 32 | P30988 |
| Tetracosactide | MC receptors | native | 24 | Q01726;Q01718;P41968;P32245;P33032 |
| Calcitonin (human) | Calcitonin receptor | native | 32 | Q16602 |
| Carperitide | NPR-A | native | 28 | P16066;P20594;P17342 |
| Bivalirudin | Thrombin | analog | 20 | P00734 |
| Nesiritide | NPR-A | native | 32 | P16066;P20594;P17342 |
| Pramlintide | Calcitonin receptor | analog | 37 | Q16602 |
| Exenatide | GLP-1 receptor | native | 39 | P43220 |
| Liraglutide | GLP-1 receptor | analog | 32 | P43220 |
| Tesamorelin | GHRH receptor | analog | 44 | Q02643 |
| Teduglutide | GLP-2 receptor | analog | 33 | O95838 |
| Lixisenatide | GLP-1 receptor | analog | 44 | P43220 |

### Training and deploying PPI predictors to score interactions

We selected 3 state-of-the-art sequence-based PPI predictors for comparison (Table 2). Two of them (PIPR and D-SCRIPT) are deep learning-based, whereas the other (SPRINT) is similarity-based.

**Table 2.** State of the art sequence-based PPI predictors

| Predictor | Type | Machine learning model | Features | Availability | Ref. |
| --- | --- | --- | --- | --- | --- |
| SPRINT | Similarity-based scoring | Not applicable | Not applicable | C++ source | 15 |
| PIPR | Machine learning | RCNN | Latent representation of protein sequences generated by a combination of convolution layers and residual gated recurrent units | Python source | 18 |
| D-SCRIPT | Machine learning | Bi-LSTM + CNN | Combination of a pre-trained embedding model with a predicted contact map | Python package | 24 |

Both PIPR and D-SCRIPT generate protein embeddings to use as inputs to deep neural networks. In other words, the protein sequences are projected into a latent space, where the coordinates relative to those of other protein pairs capture their relationship in terms of their sequence similarity or distance. There are a number of ways one may generate embeddings. The authors of PIPR^18^, for instance, concatenate the sub-embeddings obtained from residue co-occurence obtained by training a Skip-Gram model with another sub-embedding obtained by clustering amino acids into 7 classes of electrostaticity and hydrophobicity to generate a one-hot encoding (second sub-embedding). On the other hand, D-SCRIPT^24^ uses the existing Bepler & Berger embedding^25^, which is generated with a bidirectional long short-term memory neural network. The embedding generator is trained using three tasks involving two input proteins: prediction of the shared SCOP level of the proteins, self-contact maps of their 3D structures and the “soft symmetric alignment”.

While a D-SCRIPT model pre-trained on human PPIs is available, both PIPR and SPRINT require validated interaction data to make predictions. PIPR, like any machine learning-based binary classifier, requires positive and negative protein pairs for training. In contrast, SPRINT only takes interacting protein pairs (positive examples) as an input. These interactions are not used to “train” SPRINT *per se*, as SPRINT is not a machine learning method in the conventional sense, but are instead used as a database which the algorithm queries against in search of similarities with a query protein pair.

To generate the training set for PIPR and generate the SPRINT database, we retrieved all PPIs involving two human proteins from the BioGRID database (version 4.3.196)^26^ and filtered them so as to only retain high-quality interactions^27^, i.e. interactions reported by at least two groups, and detected with stringent experimental methods^27^. We added pairs assumed to be non-interacting to PIPR’s training set by adding random protein pairs to produce a balanced training set. We then trained PIPR for 100 epochs on this training set and generated the one-to-all curves for all 13 peptides. Similarly, we produced the one-to-all curves using SPRINT and the pre-trained D-SCRIPT model.

Given that all but one selected peptides target receptors expressed on the cell surface, all one-to-all curves were generated using proteins predicted to be exposed on the cell’s surface as opposed to the entire proteome. We obtained the list of the 2,886 proteins predicted to make up the human surfaceome from The *in silico* surfaceome^28^. We added thrombin, a protein involved in the coagulation cascade to all curves, because it is targeted by one of the therapeutic peptides (bivalirudin).

### Simulating peptide screening in de novo engineering

Given that all peptides under study are actually truncated segments of endogenously expressed proteins (“native” in Table 1) or recombinant analogs, training the models using the endogenous equivalents of the therapeutic peptides represents an optimistic scenario. In that “optimistic” scenario, we know that an interaction highly similar or identical to that we are attempting to predict is present in the training data. This mimics a scenario where we wish to design a peptide with some similarity to an endogenously expressed protein known to interact with the protein target.

For this reason, we ran `blastp`^29^ with the default parameters to identify proteins which share a region of high similarity with the therapeutic peptides. All interactions involving hits found via the BLAST searched were removed from our PPI dataset to produce training data for a “pessimistic” scenario representing the *de novo* case. This pessimistic scenario, denoted (-) as opposed to the optimistic scenario (+), reflects what one may encounter when designing peptides against a target for which no interactors are known.

After re-training PIPR on the pessimistic dataset, we re-generated the one-to-all curves using this pessimistic PIPR model and SPRINT.

## Results

### SPRINT outperforms deep learning models

The rank of the targeted protein(s) along the one-to-all curve conveys a lot of information with respect to the potential of a peptide as a therapeutic, since it is desirable to optimize for specific interactions. In general, a peptide whose target protein is outranked by hundreds or thousands of off-target proteins on the one-to-all curve is very unlikely to make a good candidate, under the assumption that the predictions are accurate.

Figure 3A displays the distribution of ranks of the targets on the one-to-all curve for all the therapeutic peptides considered. We observe a wide spread in the ranks of the targets along the one-to-all curves, indicating variable degrees of success in predicting a specific interaction between the therapeutic peptides and their target(s).

**Figure 3.**
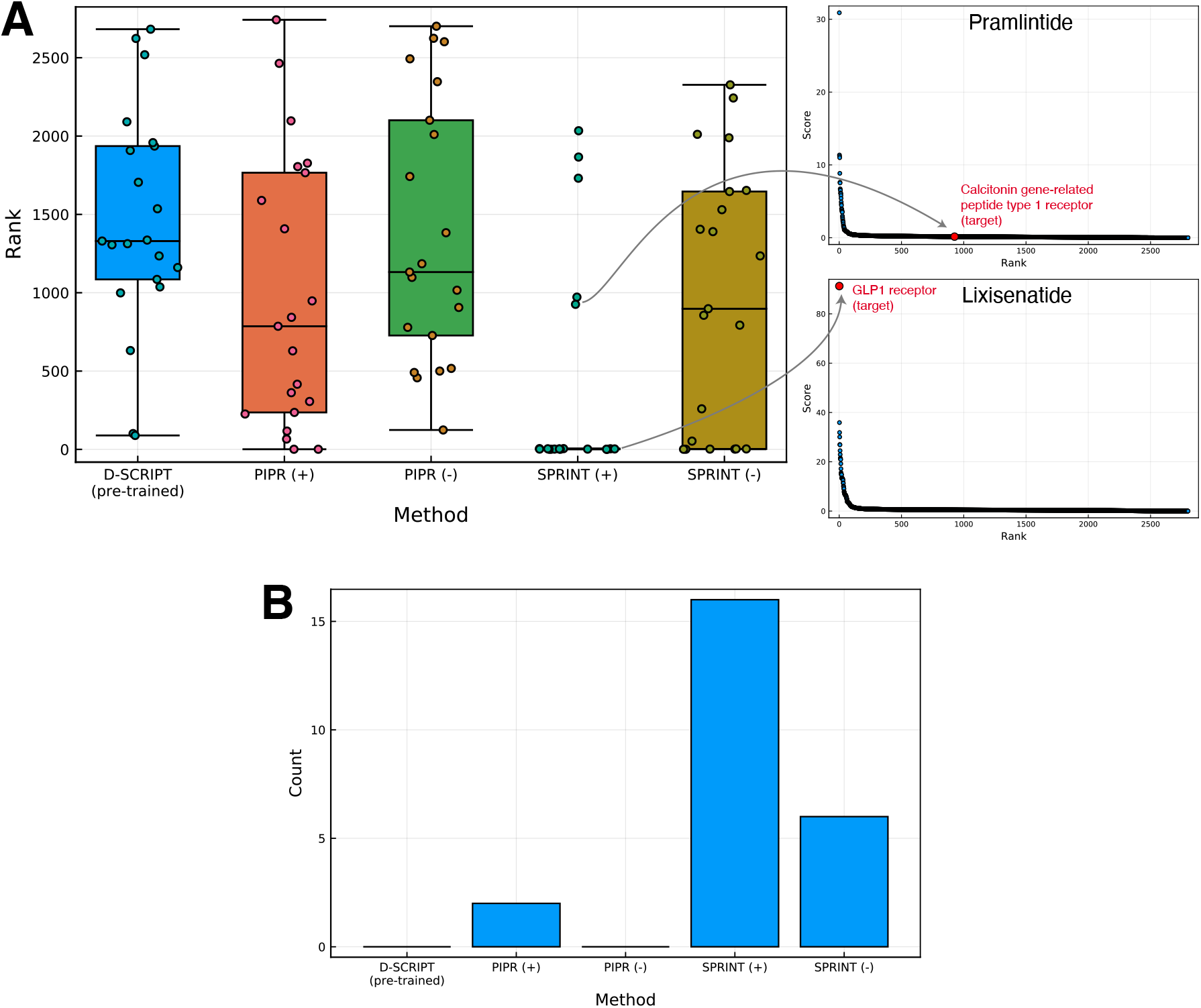
Rankings of interactions involving peptide-target pairs. (A) Distributions of ranks of the peptide-target pairs are shown for models whose training data included interactions involving an endogenous peptide analog (optimistic scenario; +) and models trained without (pessimistic scenario; -). The ranks of the true targets for two therapeutic peptides are illustrated on their one-to-all curves on the right side of the box plot. (B) Counts of therapeutic peptide targets ranking in the top 1% on the corresponding one-to-all curve.

With the aforementioned assumption in mind, we found that D-SCRIPT did not assign high ranks to the known targets on the one-to-all curves, but also failed to predict an interaction for any of the peptide-target pairs, assigning an interaction probability close to 0 in each case (see Supplementary material). The median rank of the peptide-target pairs on the respective one-to-all curves is 1,330/2,886, with only one peptide-target pair ranking in the top 500 on its corresponding one-to-all curve, and none ranking in the top 1% (Figure 3B).

PIPR performed better than D-SCRIPT, for both scenarios considered, though this difference was not statistically significant when looking at PIPR (+) (*p* = 0 06; Wilcoxon rank-sum test) and PIPR (-) (*p* = 0 60). In the pessimistic scenario analogous to assessment of *de novo* peptides, PIPR predicted an interaction (interaction probability >50%) for 3 out of the 21 peptide-target pairs, though no peptide-target pairs ranked in the top 1% on their one-to-all curve. In the optimistic scenario, however, where interactions involving proteins with high similarity to the therapeutic peptides are included in the training interactions, PIPR ranks five peptide-target pairs in the top 1% of their respective one-to-all curve. While PIPR did rank targets more highly than non-targets in more cases than D-SCRIPT, it should be noted that it produced a number of curves with a sigmoidal morphology with many high-scoring proteins (see Supplementary Figures). This suggests that PIPR may be particularly prone to false positives.

SPRINT is the predictor that achieved the highest success, scoring six true targets among the top 1% proteins on their curve in the pessimistic scenario, and 16 in the optimistic scenario. The median rank of the peptide-target pairs is also predicted to be lower by SPRINT than by PIPR in both scenarios. However, SPRINT produced statistically better ranks when comparing the methods for the optimistic scenario (+) (*p* < 0 01), but not for the pessimistic scenario (-) (*p* = 0.13). In general, it appears that SPRINT outperforms deep learning predictors for the task of predicting protein-peptide interactions, particularly in the optimistic scenario where all known interactions are leveraged to make predictions.

### Sensitivity to point mutations

We found that SPRINT and PIPR, in the optimistic training scenario, were able to detect point mutations in the sequence of the peptides, by computing the one-to-all curves for all mutants produced by mutating every single non-Gly amino acid in the sequence to Gly (Figure 4). Both methods produced positive and negative changes in score upon mutation. However, these changes were less likely to result in a change in the target ranks for SPRINT than PIPR, because the target scores were often well above that of non-target proteins on SPRINT-generated curves, and this gap was rarely offset by a drop in score.

**Figure 4.**
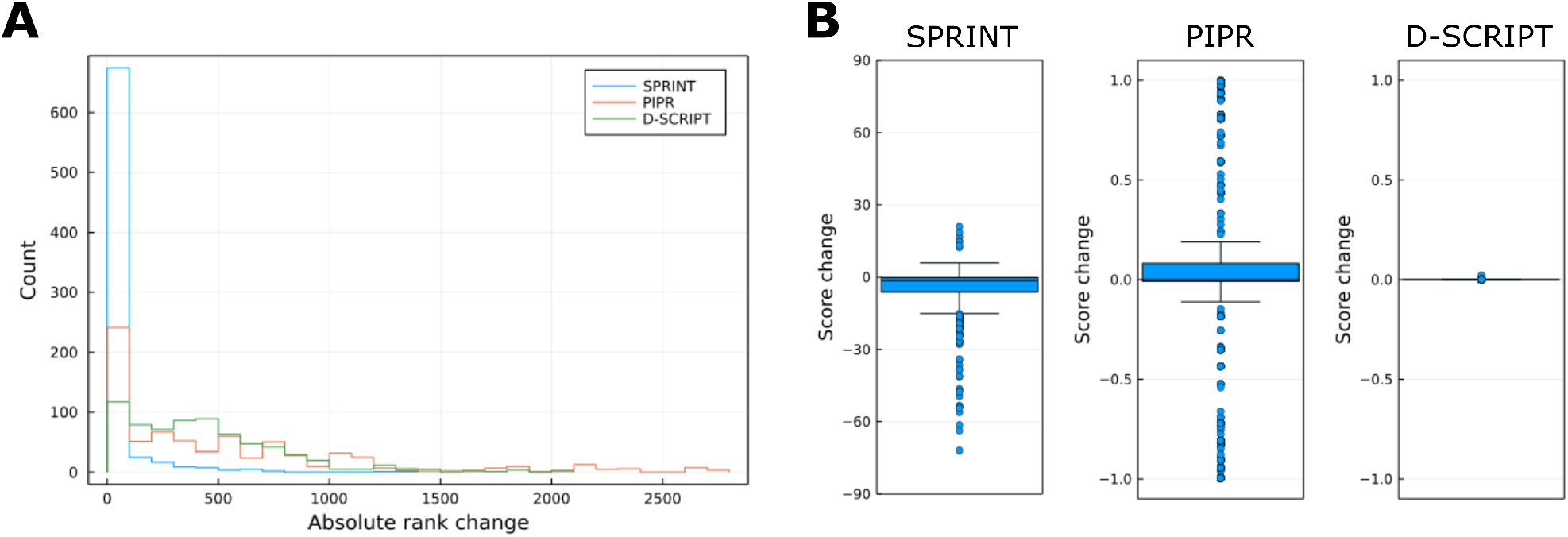
Distribution of changes in rank and score. The one-to-all curves were regenerated for each peptide by mutating each non-Gly residue to Gly to evaluate (A) the absolute change in rank on the one-to-all curve and (B) the change in score. As a reminder, SPRINT scores are unbounded, while PIPR and D-SCRIPT scores are probabilities that range from 0 to 1.

Even though mutations led D-SCRIPT to produce large rank changes, those are an artifact of the position of the targets in the baseline portion of the one-to-all curve. Small changes in score can produce very large changes in rank when the scores are very close to 0, as is the case for target scores generated by D-SCRIPT. Looking only at the scores produced by D-SCRIPT, it appears that it is rather insensitive to point mutations in the peptide sequence.

### Protein-protein interaction predictors are useful for peptide analog screening

Consistent with the results above, screening for peptide analogs with some similarity to endogenously expressed proteins is a scenario where SPRINT outperforms the other predictors, as illustrated with lixisenatide in Figure 5.

**Figure 5.**
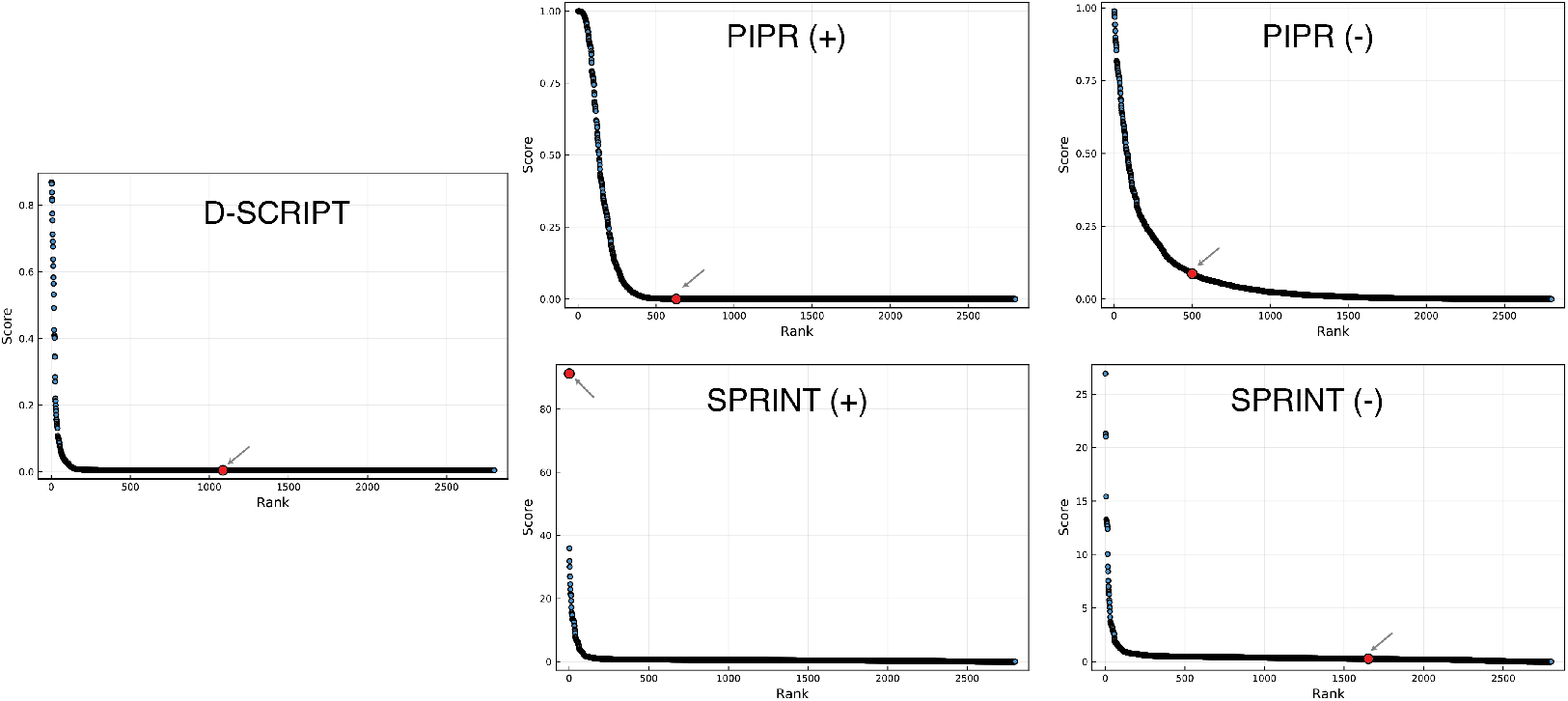
One-to-all curves for lixisenatide. The one-to-all curves generated using the predicted interaction scores of the three predictors are shown here. The true target protein is highlighted in red, while the black dots represent off-target proteins. One-to-all curves generated with predictors trained in the optimistic scenario are indicated with a (+).

Lixisenatide is a 44 amino acids glucagon-like peptide-1 agonist marketed by Sanofi and used in the treatment of type 2 diabetes to increase insulin secretion. This peptide is an GLP-1 *analog* corresponding to a portion of the exendin-4 protein found in the venom of Gila monster (*Heloderma suspectum*)^30^. The target, the glucagon-like peptide 1 receptor (Accession ID: P43220), is a G-protein coupled receptor located on the cell surface.

The one-to-all curves show that both D-SCRIPT and PIPR assign low interaction probabilities between lixisenatide and the GLP1 receptor. The target proteins rank poorly along the curves. In fact, the probabilities of interaction are well below 0.5, thus no interactions are predicted to occur. In the case of D-SCRIPT, 16 proteins are predicted to interact with lixisenatide, none of them being the GLP1 receptor. In fact, D-SCRIPT predicts a 0.4% chance of an interaction between lixisenatide and the GLP1 receptor. PIPR predicts an interaction with higher probability, though it still fails to meet the 0.5 threshold while assigning higher scores to over 500 non-target proteins on the cell’s surface in both the optimistic and pessimistic scenarios. SPRINT, when “trained” in the optimistic scenario, accurately ranks the GLP1 receptor as first on the one-to-all curve while assigning much lower scores to other off-target proteins.

Only SPRINT makes credible predictions under the assumption that lixisenatide interacts specifically with the GLP1 receptor, and only in the optimistic case, i.e. when the interaction between (pro)-glucagon (Accession ID: P01275) and the GLP1 receptor is included in the training data. In fact, alignment shows that lixisenatide, though not derived from human glucagon, aligns to a certain degree with a portion of it (98-141), part of which (98-128) is normally cleaved to yield the GLP1 peptide Figure 6. This suggests that SPRINT can effectively be leveraged to screen peptide analogs that are somewhat similar, but not identical to endogenously expressed proteins.

**Figure 6.**
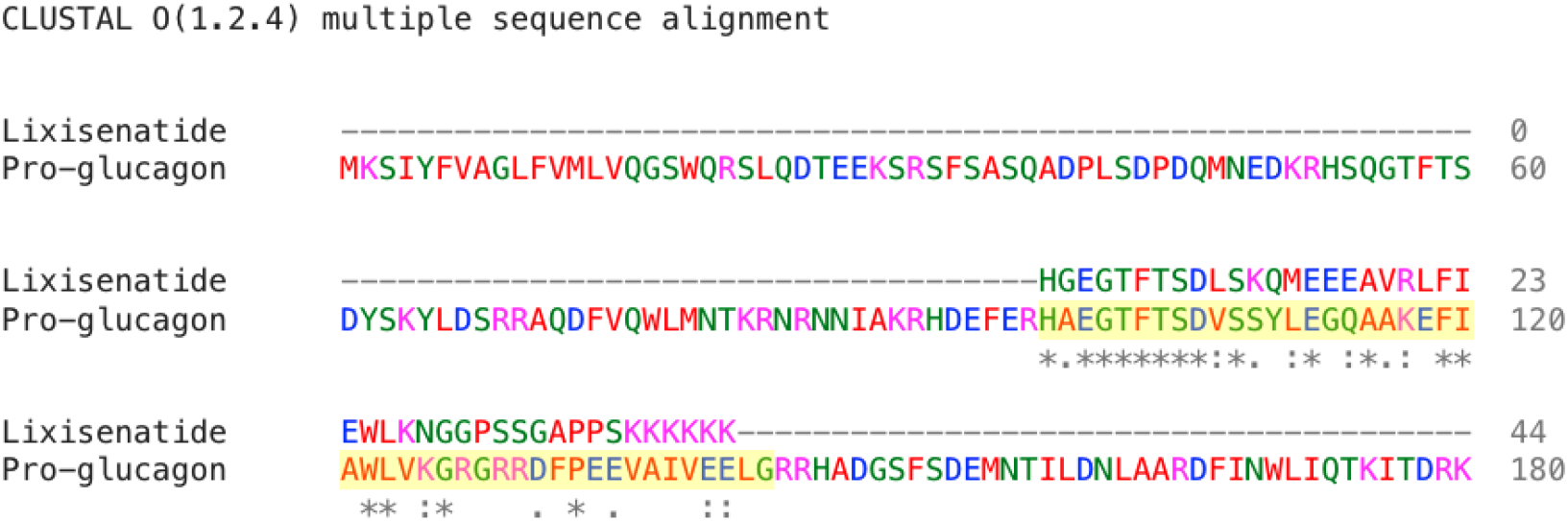
Alignment between lixisenatide and pro-glucagon. Results of the alignment between lixisenatide and pro-glucagon generated with Clustal Omega^31^. The highlighted region corresponds to the endogenously expressed GLP1 peptide.

### Assessing peptide fitness for de novo design remains challenging

All predictors, including SPRINT, generally failed to rank targets highly on the one-to-all curve in the more challenging *de novo* screening scenario, i.e. where interactions involving proteins with high similarity with to the therapeutic peptides were excluded from the training data.

In a limited number of cases, SPRINT assigns better ranks to peptides in the pessimistic scenario. Such was the case for carperitide, a peptide marketed by Daiichi Sankyo for the treatment of acute heart failure. Both in the presence and absence of training interactions involving the *α*-atrial natriuretic peptide, of which carperitide is a recombinant version, SPRINT ranked the intended target highly on the one-to-all curves (Figure 7), albeit with a lower score than in the optimistic scenario (+). In these curves, all three best-ranked proteins are isoforms of the atrial natriuretic peptide receptor. We observed similar results for nesiritide, which is also a recombinant version of a natriuretic peptide.

**Figure 7.**
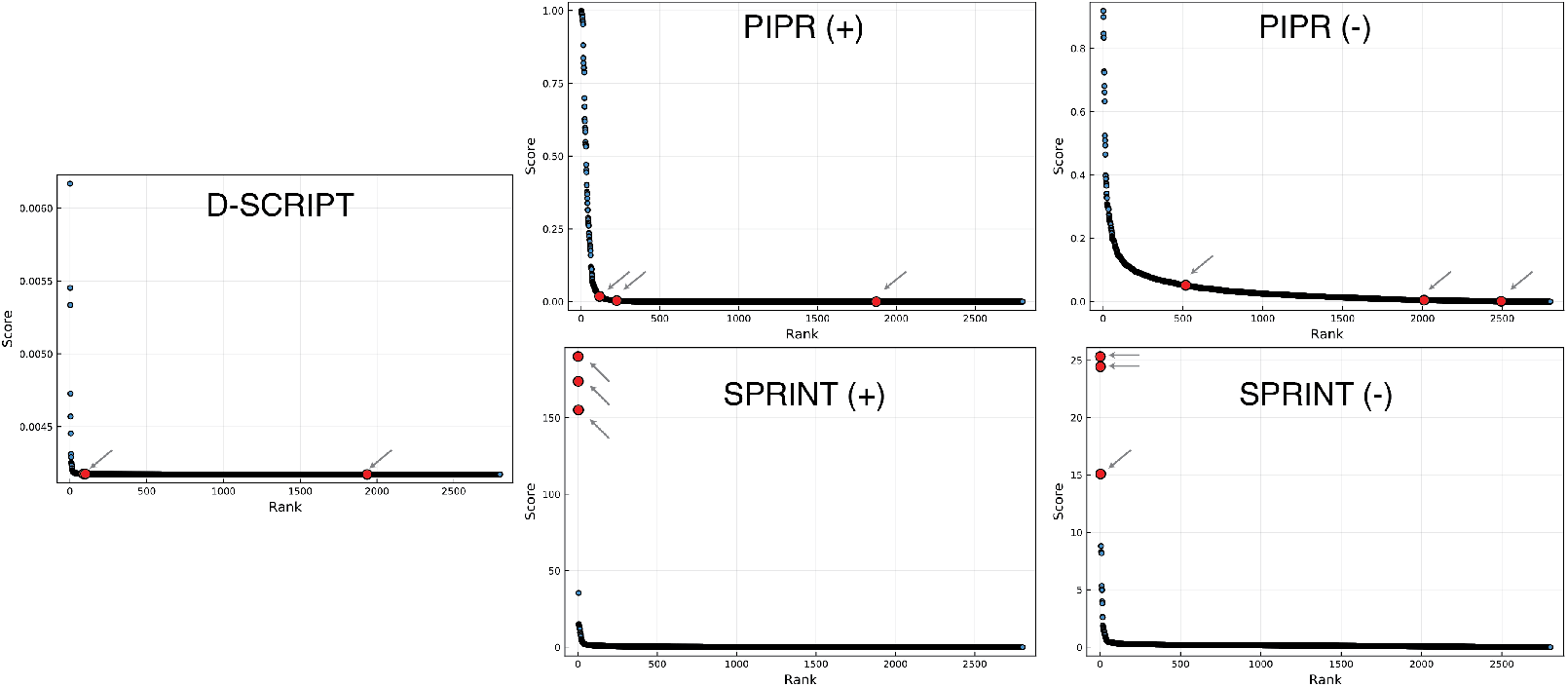
One-to-all curves for carperitide. The one-to-all curves generated using the predicted interaction scores of the three predictors are shown here. The true target protein is highlighted in red, while the black dots represent off-target proteins. One-to-all curves generated with predictors trained in the optimistic scenario are indicated with a (+).

## SPRINT and the GLP receptors

Given that the Glucagon-like peptide receptors 1 and 2 share high similarity, we expected both of them to rank similarly on the one-to-all curves of GLP1R and GLP2R agonists. We observed, using SPRINT (+), that GLP1R was ranked first for all GLP receptor-targeting peptides (Figure 8). This was the case even for Teduglutide which is a glucagon-like peptide 2 analog. In all cases, GLP2R is outranked by Glucagon receptor and the Gastric inhibitory polypeptide receptor with which it aligns well (identity > 40%).

**Figure 8.**
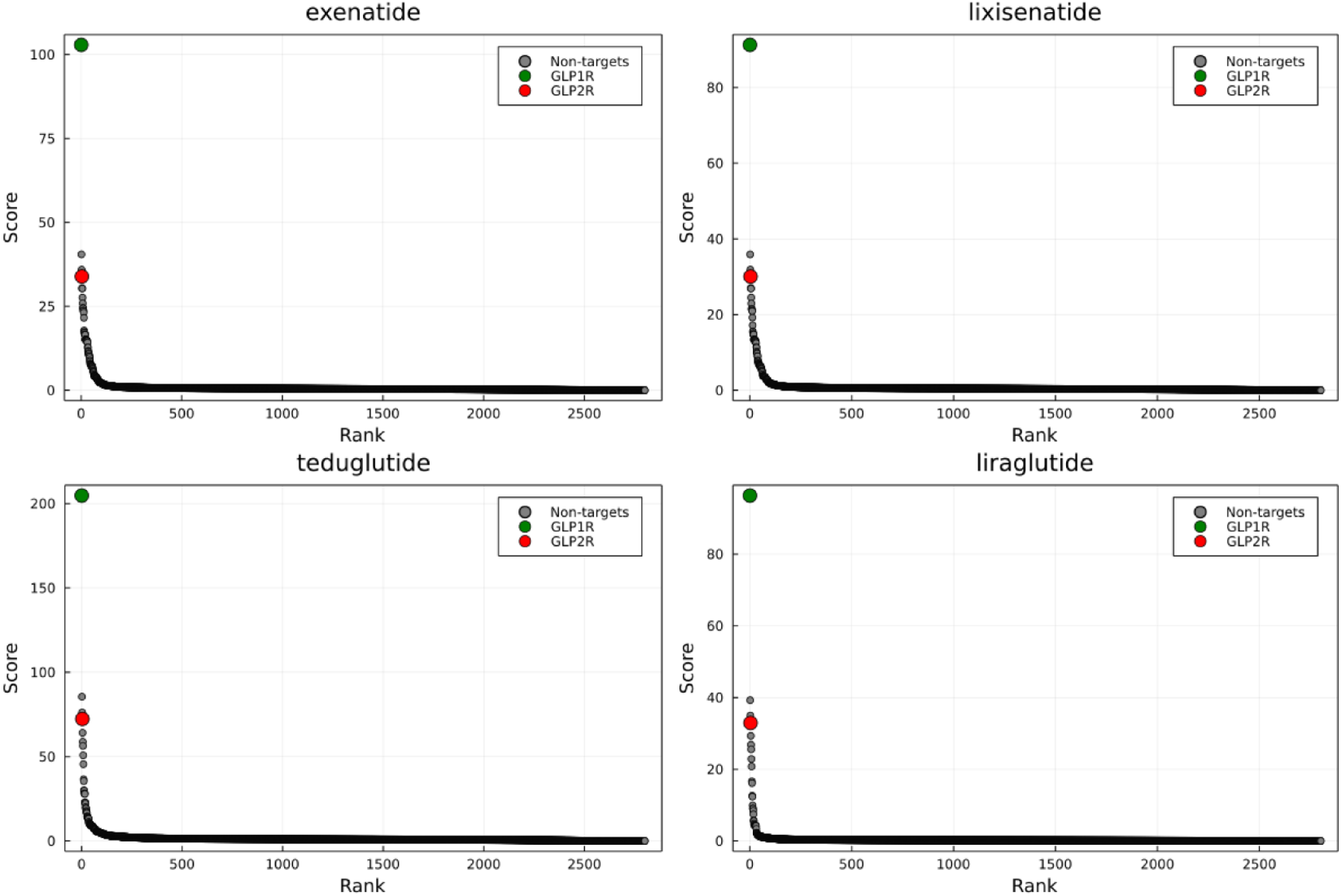
One-to-all curves of the peptides targeting a GLP receptor. Exenatide, liraglutide and lixsenatide target the GLP-1 receptor while teduglutide was developed to target the GLP-2 receptor. The curves were generated with SPRINT (+).

We also applied SPRINT to a novel chimeric peptide built from GLP-2 that replaces the C-terminus end of the peptide with that of GLP-1 (residues 20-33) published in 2020^32^. The authors demonstrated that the peptide activate the GLP1R and GLP2R roughly equally. In spite of this, SPRINT (+) ranked GLP1R 2nd, and GLP2R 13th, though with an appreciable difference in score of 116.4 (Figure 9). However, it is worth pointing out that both receptors ranks highly and are well above the knee of the one-to-all curve. Oddly, GLP1R is outranked by a Tumor necrosis factor ligand (accession id: P48023) that shares limited identity (23%) with GLP1R. In all cases, the one-to-all curve suggests that this chimeric peptide may be less specific than the other GLP analogs assessed in the current study.

**Figure 9.**
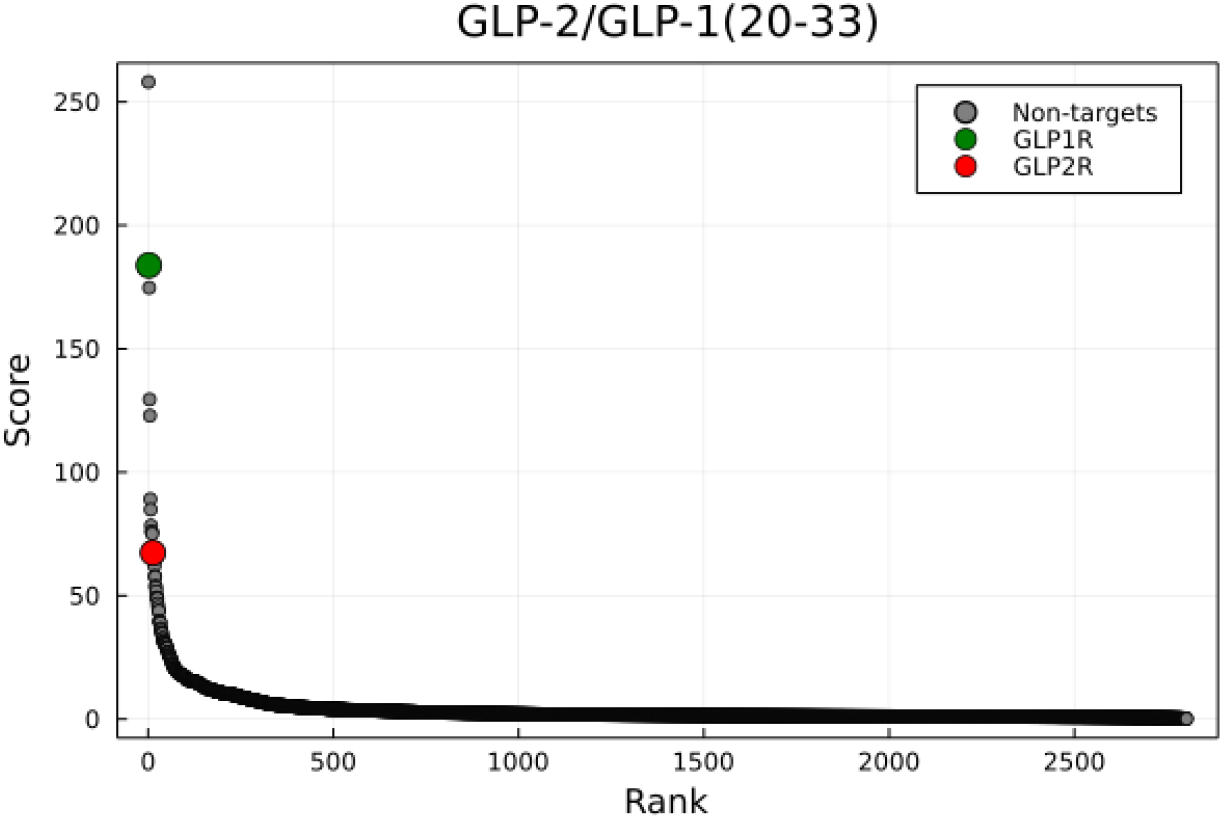
One-to-all curve of a chimeric peptide with roughly equal affinity for GLP1R and GLP2R. The one-to-all curve of the chimeric peptide GLP-2/GLP-1(20-33) generated with SPRINT (+).

## Discussion

The ability to rapidly and inexpensively screen peptides *in silico* as potential therapeutics targeting a specific protein is highly desirable, but remains challenging. This can be partially attributed to the relative scarcity of validated peptide-protein interactions. Given that most PPI predictors are trained using long proteins, we anticipated that these methods would perform poorly when deployed for the task of predicting interactions involving shorter peptides (<50 amino acids).

### The case for similarity-based methods

D-SCRIPT and PIPR, both deep learning methods, mostly failed to produce realistic one-to-all curves. However, SPRINT produced reasonably good one-to-all curves that ranked known targets better than non-targets, particularly when all available protein-protein interactions were used to make predictions. This suggests that SPRINT may be suitable as a tool to screen peptides in cases where an endogenous interactor is known for the target protein. This explains why curves for GLP analogs produced more realistic results than other peptides such as peptides targeting the calcitonin receptor (calcitonin and pramlintide). The interaction data used for the experiments does not contain proteins with high similarity to calcitonin, whereas it did contain interactions involving glucagon and GLP receptors. This further highlights the importance of the training data. Deep learning methods, in contrast, are generally not suitable. We reason that SPRINT outperforms the competing methods because of its prediction mechanism. Deep learning methods make predictions based on both the global and local properties of protein sequences which are captured in an embedding. This is in contrast with SPRINT, whose predictions rely on small, local protein regions of similarity that are roughly the length of a peptide (~20 amino acids). The fact SPRINT uses what essentially amounts to short peptides as evidence to support their predictions makes SPRINT and other similarity-based PPI predictors (eg. PIPE^16^) uniquely suited for such tasks. We also found that point mutations in the peptide sequence could be detected by SPRINT, though they could also be detected by PIPR. This is interesting, as it implies that the methods may have sufficiently high resolution to tune peptides at the residue scale, though more work would be needed to determine how useful it is in practice.

### Sequence-based methods achieve limited success in de novo design tasks

Unsurprisingly, we found that assessing the quality of peptides that have little to no high-similarity regions to proteins in the training dataset of interacting proteins is challenging. In fact, SPRINT is sensitive to small changes in the training data, as top-ranking peptide-target pairs often dropped to the bottom of the one-to-all curve when one or two key interactions were removed from the training data (pessimistic scenario). For this reason, applying SPRINT to screen *de novo* peptides for potential interactions with a target is likely to be unsuccessful, depending on the training data. However, SPRINT assigned high ranks to the target (natriuretic peptide receptor A) for both carperitide and nesiritide, even in the absence of their endogenous equivalent in the training data under the pessimistic scenario. Our data do not support the generalization of this result to other peptides, especially considering that carperitide and nesiritide target the same receptor.

### Rank on the one-to-all curve is a key metric

SPRINT produces an unbounded sum of the number of regions of similarity between query proteins and proteins in the training set of interacting proteins, weighted by the similarity as scored with a substitution matrix. That raw sum corresponds to the prediction score, and the threshold separating non-interacting pairs from interacting pairs is arbitrary. Normally, a reasonable threshold could be determined through cross-validation for normal protein-protein interaction tasks. We believe the rank along the one-to-all curve to be just important than the raw score of a target, especially in the context of peptide engineering. A low predicted interaction score may be due to limited similarity to regions in known interacting proteins. However, for peptide engineering tasks, such a cross-validation experiment is not possible, since no a novel peptide would have no known and validated interactions. Thus it may be worth validating a high-ranking peptide *in vitro*, even if the raw interaction score is low. More work will be needed to develop a peptide fitness metric that incorporates both the rank of the target protein along the one-to-all curve, and the predicted score and other features derivable from the curve. The curves generated for GLP receptor agonists with SPRINT also support the idea that rank is just as, if not more, insightful than the prediction score. We saw that although Teduglutide targets GLP2R and GLP1R was ranked higher, GLP2R still ranked very high on the one-to-all curve. We observed similar results for a chimeric peptide demonstrated to activate the GLP1R and GLP2R roughly equally. It is worth noting that the prediction scores are meant to reflect the degree of confidence in an interaction, not the binding affinity, which is a different task.

### Tackling the challenges associated with de novo peptide design

While general-purpose methods like SPRINT show some potential in a purely *in silico* context for design of peptide analogs, we believe that incorporating target-specific training data obtained experimentally could increase the applicability of similarity-based PPI prediction methods for *de novo* peptide engineering tasks. For example, sparsely sampling the peptide space (~1,000 peptides) and testing for interactions with the target protein using peptide arrays could supplement the initial training data and drastically improve the predictions. In fact, one group found that sparse sampling of the peptide space for target-specific peptides could provide the training data necessary to train a simple neural network that could predict with reasonable accuracy whether a peptide would bind to that specific target^33^. Using predictors like SPRINT in an adaptive fashion could also significantly enhance their applicability in peptide engineering task. For instance, peptides optimized *in silico* could periodically be tested experimentally for binding to the target. Hits could then be incorporated in the training data in an iterative fashion and improve the quality of the predictions in future iterations.

### Limitations

At the time of writing this paper, few peptides marketed for therapeutic use met the criteria for inclusion in this study. Furthermore, the peptides included in this study targeted only a small set of receptors. Therefore, more work will be needed to determine whether these results hold up when applied to a more diverse set of protein targets.

In addition, given the extremely rapid pace of discovery in deep learning, it is not unreasonable to expect that deep learning methods outperforming similarity based methods may be developed in the near future. We are aware of two novel methods published while this article was undergoing review. CAMP is a novel deep learning method that has shown promise in assessing not only protein-peptide interaction probabilities, but also in identifying the peptide residues that underpin the interaction using convolutional networks and a self-attention mechanism^34^. Mutual Information Maximization Meta-Learning is another approach relying on meta-learning and information theory that optimizes peptide bioactivity^35^. In fact, the growing interest in computational therapeutic peptide screening may require us to regularly benchmark emerging methods.

### Concluding remarks

In summary, our work shows that similarity-based PPI predictors are currently more suitable than deep learning methods to evaluate the potential of peptide therapeutics. We also showed that deploying such methods for *analog* peptide engineering is more likely to be successful than for engineering peptides *de novo*, i.e. against targets for which no interactors are known. We anticipate that more specialized protein-peptide interaction predictors will become available as more data becomes available, but believe that existing methods can already be integrated in peptide engineering pipelines. Finally, we believe that adaptive use of existing sequence-based PPI prediction methods holds great potential.

## Acknowledgements

The authors would like to thank the Natural Sciences and Engineering Research Council of Canada (NSERC) who provided financial support to F.C. through a PGS-D3 award.

## Author contributions statement

F.C. conceived and conducted the experiment(s), F.C., K.K.B. and J.R.G. analyzed the results. F.C. wrote the manuscript. K.K.B. and J.R.G. edited the manuscript. All authors reviewed and approve of the manuscript.

