## Supplemental material for "Assessing sequence-based protein-protein interaction predictors for use in therapeutic peptide engineering"

### Supplementary figures

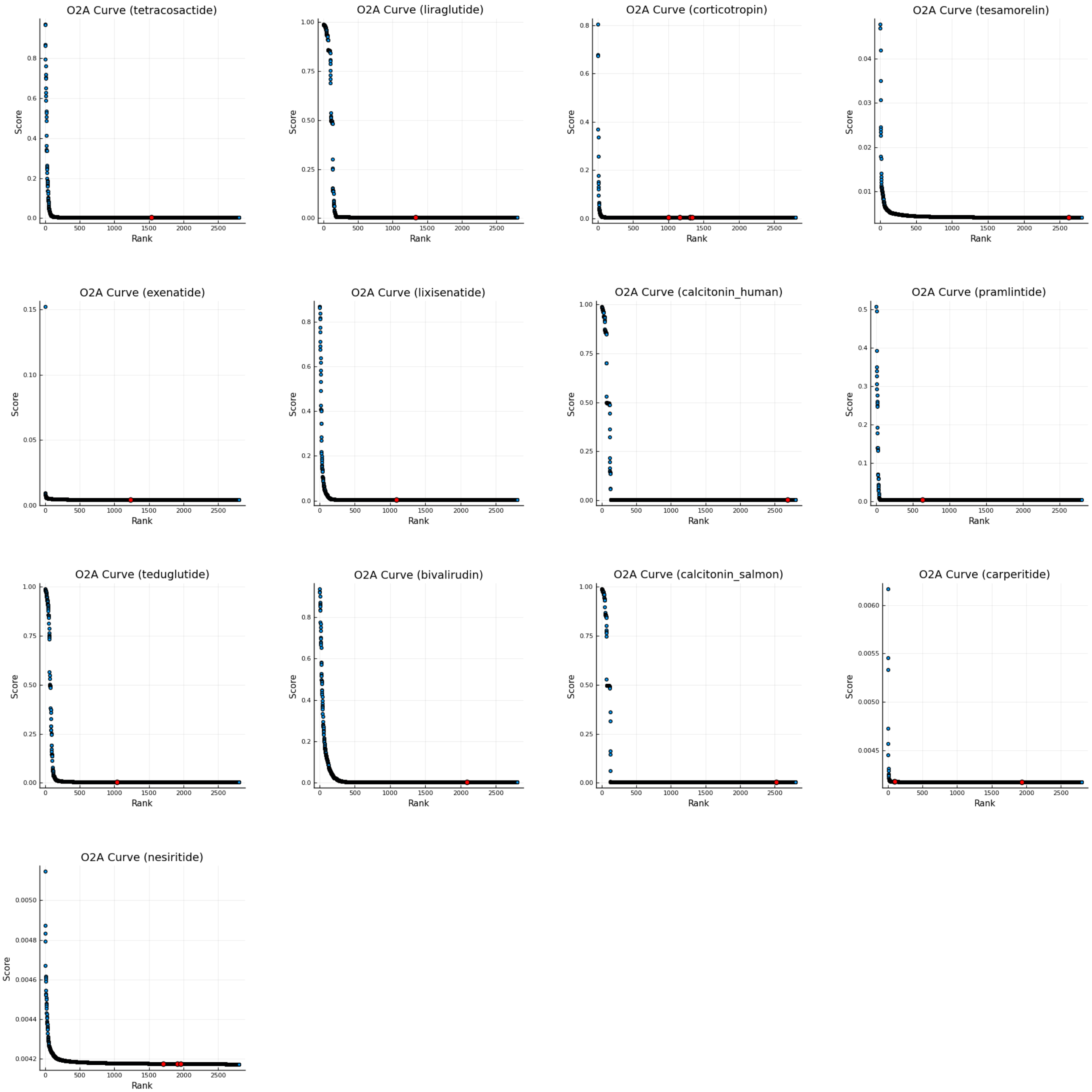


**Supplementary Figure 1.** **All one-to-all curves generated with D-SCRIPT.**

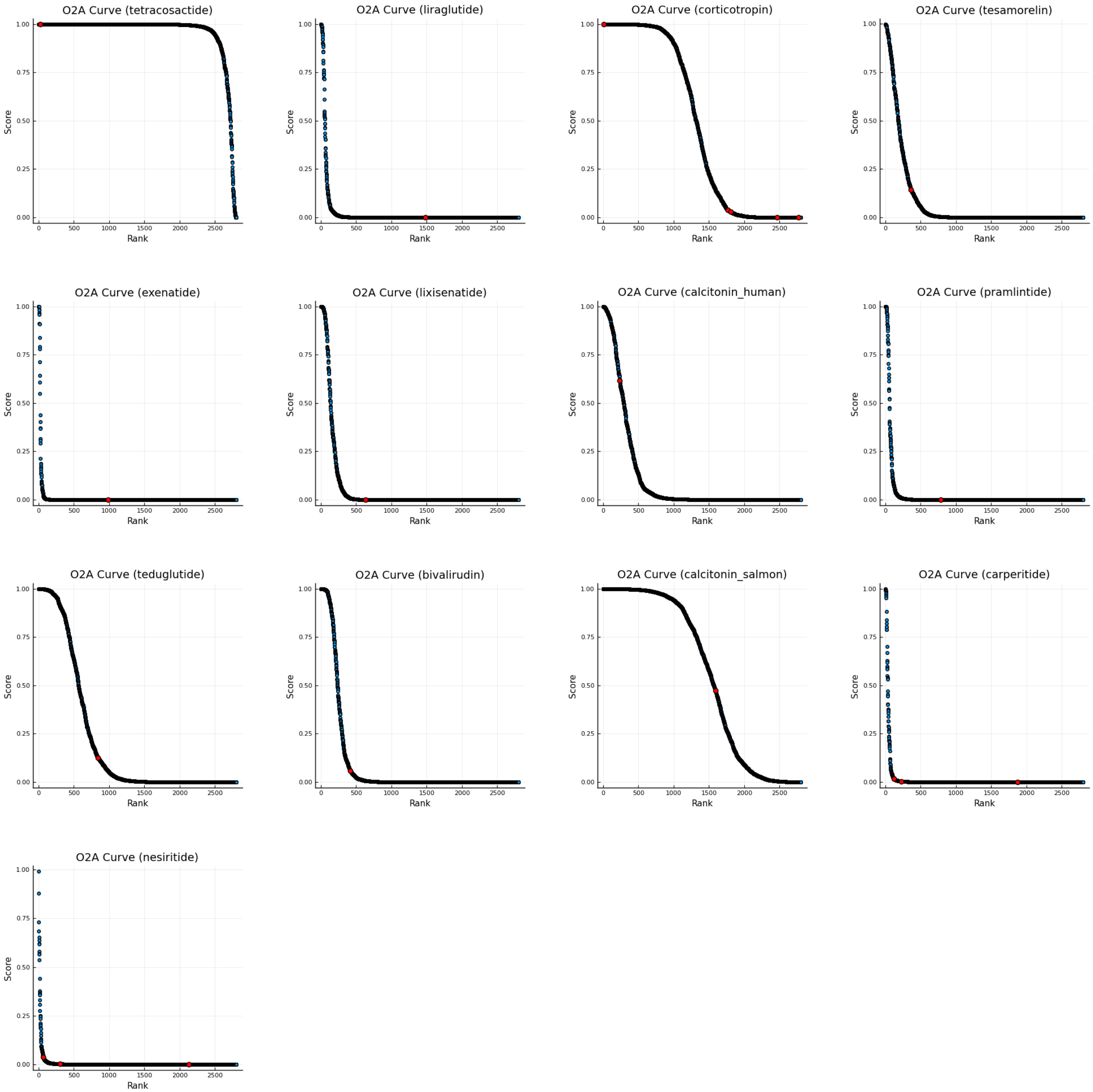


**Supplementary Figure 2.** **All one-to-all curves generated with PIPR (optimistic scenario).**

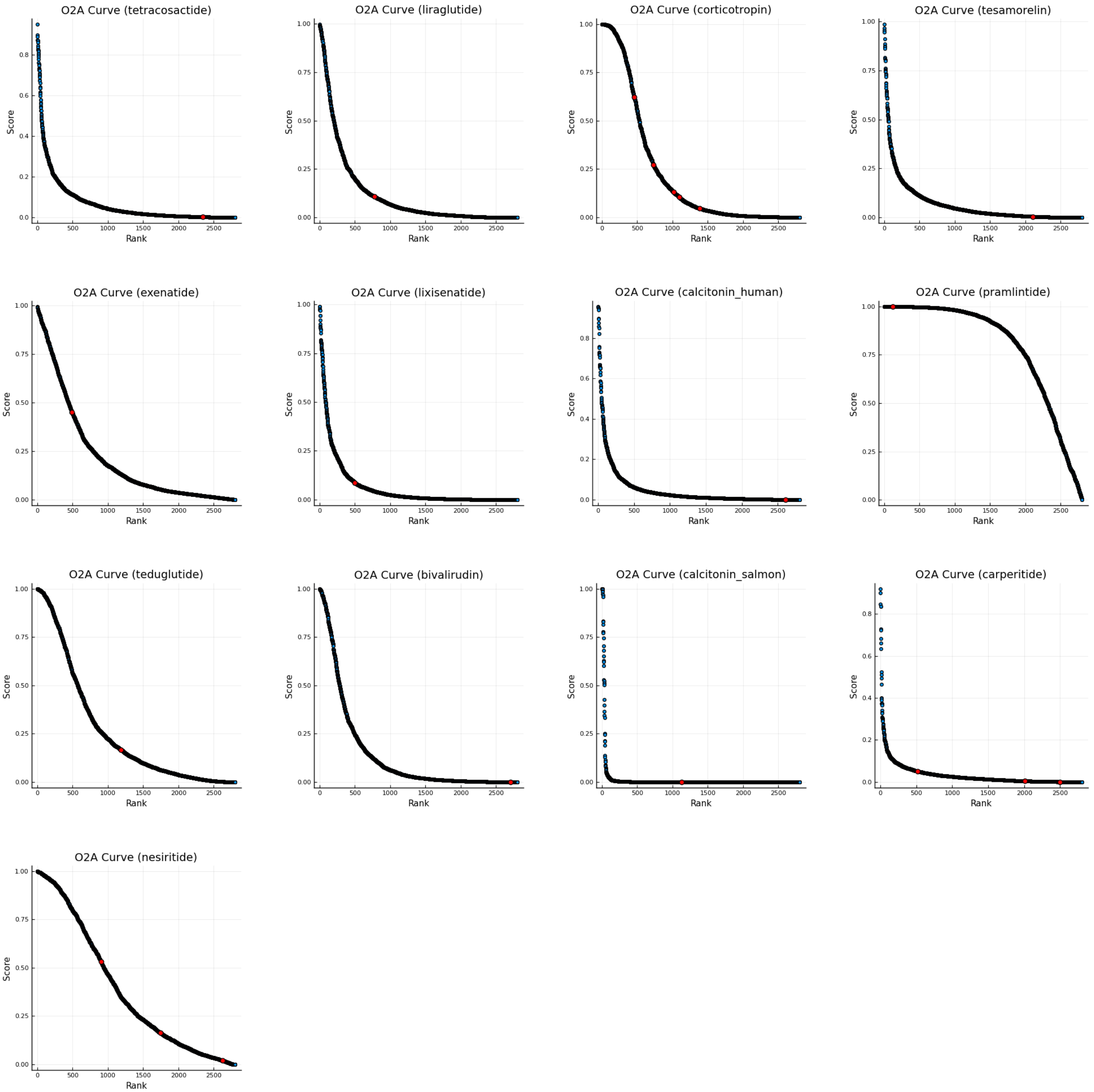


**Supplementary Figure 3.** **All one-to-all curves generated with PIPR (pessimistic scenario).**

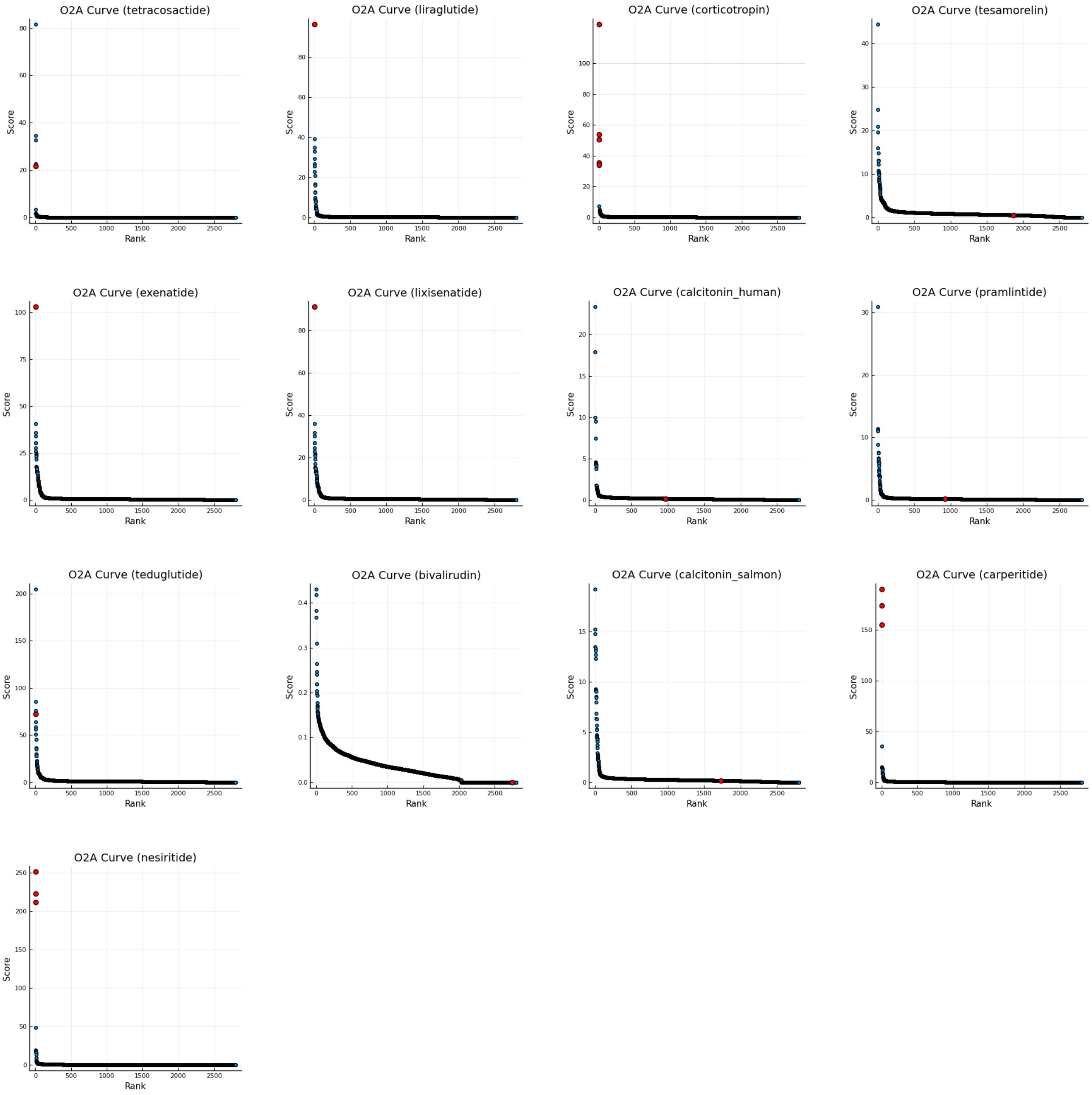


**Supplementary Figure 4.** **All one-to-all curves generated with SPRINT (optimistic scenario).**

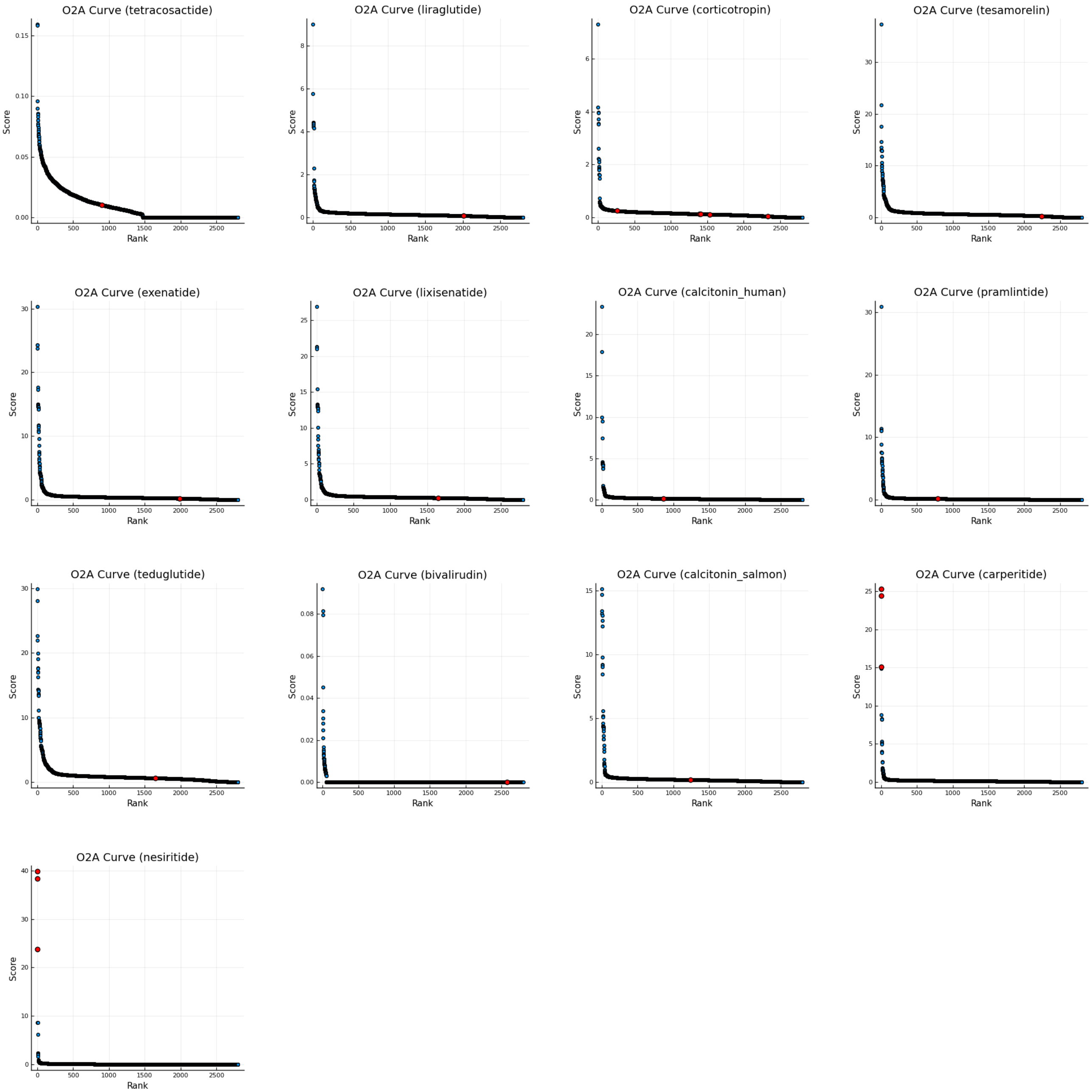


**Supplementary Figure 5.** **All one-to-all curves generated with SPRINT (pessimistic scenario).**
